# Bund removal to re-establish tidal flow, remove aquatic weeds and restore coastal wetland services – North Queensland, Australia

**DOI:** 10.1101/639799

**Authors:** Brett N. Abbott, Jim Wallace, David M. Nicholas, Fazlul Karim, Nathan J. Waltham

**Author notes:** Corresponding author –.

## Abstract

The shallow tidal and freshwater coastal wetlands adjacent to the Great Barrier Reef lagoon provide a vital nursery and feeding complex that supports the life cycles of marine and freshwater fish, important native vegetation and vital bird habitat. Urban and agricultural development threaten these wetlands, with many of the coastal wetlands becoming lost or changed due to the construction of artificial barriers (e.g. bunds, roads, culverts and floodgates). Infestation by weeds has become a major issue within many of the wetlands that were modified (bunded) for ponded pasture growth last century. A range of expensive chemical and mechanical control methods has been used to try and restore some of these coastal wetlands, with limited success. This study describes an alternative approach to those methods, investigating the impact of tidal reinstatement after bund removal on weed infestation, associated changes in water quality, and fish biodiversity, in the Boolgooroo lagoon region of the Mungalla wetlands, East of Ingham in North Queensland. High resolution remote sensing, electrofishing and in-water logging was used to track changes over time – 1 year before and 4 years after removal of an earth bund. With tides only penetrating the wetland a few times yearly, gross changes towards a more natural system occurred within a relatively short timeframe, leading to a reduction in weed infestation, reappearance of native vegetation, improvements in water quality, and a tripling of fish diversity. Weed abundance and water quality does appear to oscillate however, dependent on summer rainfall, as changes in hydraulic pressure stops or allows tidal ingress (fresh/saline cycling). With an estimated 30% of coastal wetlands bunded in the Great Barrier Reef region, a passive remediation method such as reintroduction of tidal flow by removal of an earth bund or levee could provide a more cost effective and sustainable means of controlling freshwater weeds and improving coastal water quality into the future.

## Introduction

Coastal floodplains around the world have been modified for human gain, most notably being hydrologically altered either totally or partially reducing connectivity between floodplains and coastal areas [1, 2]. Floodplain, coastal tidal and freshwater wetlands are important habitat because they provide important biodiversity, hydrological, cultural and economic goods and services [3–5]. However, these wetlands are under great pressure due to urban and industrial development [6, 7] or agricultural and grazing land expansion, with many coastal wetlands becoming lost due to the construction of artificial barriers (e.g. bunds, roads, culverts and floodgates) which have stopped or reduced tidal flushing, negatively impacting aesthetic and ecological values [8]. The widespread degradation of coastal wetlands has led to major shifts in species assemblages and declines in aquatic species productivity. In response, there has been increased effort to rehabilitate coastal wetlands by removing these artificial barriers [9, 10], and provide protection and restoration of coastal wetlands [11] to improve their ecosystem services including connectivity and functionality as productive fish habitat [12, 13] and also deliver opportunities for carbon sequestration and storage [14].

The coastal wetlands of north Queensland contain unique and valuable biodiversity at the interface between two World Heritage areas; the Great Barrier Reef and Australia’s tropical rainforests [15, 16]. These wetlands are important ecological assets with significant cultural and economic values [17]. In their natural state they provide habitat for native plants and animals, migratory birds as well as potential water quality improvement and hydrological regulation functions [18–20]. The aboriginal peoples of Australia associate great cultural value to wetlands [21, 22], which also have commercial and recreational value [23, 24]. However, many of the wetlands along the north Queensland coast, and the services they provide, have been lost; for example, it has been estimated that between 60 and 90% of freshwater and saline wetlands have vanished in favour of rural and urban development [6, 25, 26]. Of the wetlands that remain, many are degraded via a combination of earth bunding, to exclude seawater and reclaim land for pasture [27–29], upstream agricultural use (grazing and sugar cane production) which leaches ecologically damaging nutrients and sediments [30, 31], and extensive aquatic invasive weed chokes [32, 33] which contributes to hypoxic conditions and fish kills [34, 35].

The Mungalla wetlands east of Ingham, on the north Queensland coast, are characteristic of the many degraded intertidal wetlands adjacent to the Great Barrier Reef lagoon [32]. The degradation began after an earth bund was constructed in the mid-1940s, initially to provide access across the wetlands [36], which excluded seawater and created a ponded pasture for grazing [21]. A short time later para grass *(Urochloa mutica)* was introduced into the freshwater ponded area, which formed above the bund. Para grass was introduced into Queensland in 1884 for improving river bank stabilization [37] and has been used as ponded pasture in bunded areas of coastal marine plains since artificial ponding for grazing started in Queensland in the 1930’s [38]. More recently two other ponded pasture species have been introduced – Olive Hymenachne *(Hymenachne amplexicaulis)* and Aleman grass *(Echinochloa polystachya).* These two species were introduced into Queensland in 1988 [39] despite apparent warning of possible weedy invasion by the Queensland Environmental Protection Agency [40, 41]. Subsequently the two grasses have been recognized as weeds regionally, with Olive Hymenachne being declared a weed of national significance (WONS) 10 years after release [42]. Aside from the pasture introductions, two other nationally declared weeds are present in the wetlands, Salvinia *(Salvinia molesta)*, and Water hyacinth *(Eichhornia crassipes)*, both of which are spread by spore/seed or asexually through fragmentation. Weed growth and expansion into the wetland was exacerbated by Palm Creek carrying nutrients downstream from a large area of sugar cane and a sugar mill [32]. In response, the Mungalla Wetland Management Strategy [43] prescribed weed control using aerial and ground based application of chemical herbicides in the first instance with the possibility of exploring bund removal. The initial efforts were expensive and ecologically undesirable as freshwater weed removal was partial and short-lived. With the limited success of the chemical weed control, it was decided to investigate bund removal as a natural form of weed control using tidal ingress from the seaward side of the wetland. There was however, some uncertainty around the frequency, duration and extent of seawater penetration into the wetland once the bund was removed. Hydro-dynamic modelling simulations by Karim *et al.*, [44] predicted that large (king) tides occurring in December/January and June/July each year should penetrate upstream of the existing bund. However, these simulations took no account of the impact of different depths of standing water within the wetland, which could form a hydrological barrier to, and potentially inhibit, on-going ingress of seawater. Despite these uncertainties it was decided to remove sections of the earth bund wall.

This paper gives an overview of the changes in weed infestation, fish biodiversity and detailed monitoring of the depth and quality of the water within the wetland for 1 year before, and 4 years following reinstatement of tidal flow into the Boolgooroo lagoon region of the Mungalla wetlands complex, North Queensland (Fig 1). We assess the success of the intervention in the context of the local (Mungalla) wetland management strategy [43] which specifically targeted weed removal, and explore the broader implications of this management strategy for coastal wetland restoration.

**Fig1.**
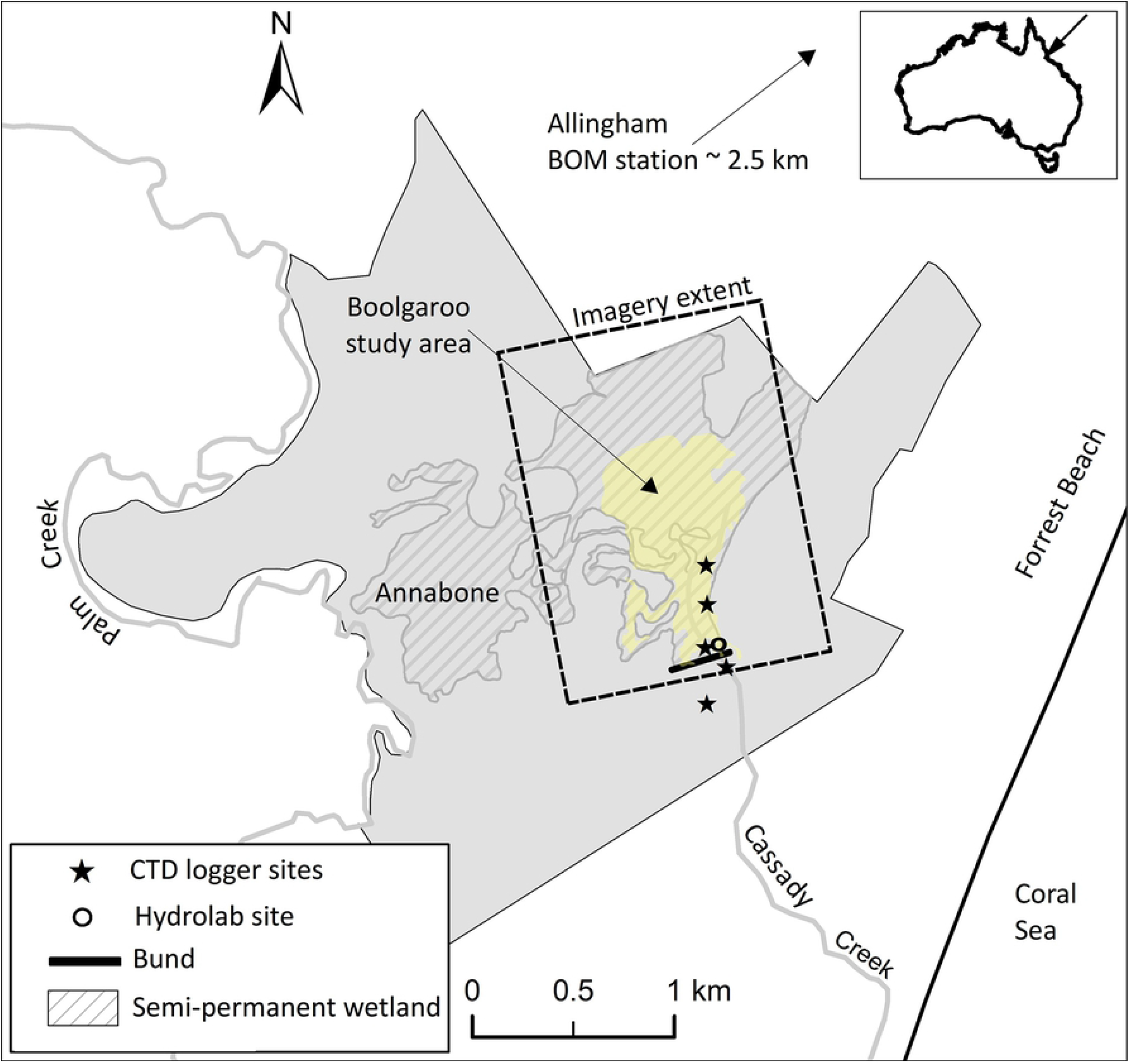
Location of wetlands within Mungalla station in the lower Herbert River catchment, Queensland, Northern Australia. The Mungalla wetlands are shown in light grey, the Boolgooroo region of the wetland complex is shown in yellow. Also shown are the locations of the logger sites above and below the earth bund which was removed on 6^th^ October 2013.

## Methods

### Location and climate

Mungalla Station (146°16’19”E, 18°42’55”S), is a 830 ha property located in the lower part of the Herbert River catchment south east of Ingham, North Queensland (Fig 1). Mungalla station has been run as a cattle-grazing enterprise for over a century, and since 1999 by the Nywaigi Aboriginal people, who also manage the wetland area. Until its removal on 6th October 2013 the wetland area above the bund was palustrine with saline estuarine wetland and saltmarsh below it. Typically saltmarsh would merge into intertidal grass-sedge wetlands dominated by Bulkuru *(Eleocharis dulcis)* [45] – the common name Bulkuru originally derives from the indigenous Boolgooroo which is a clear indicator of the wetland’s pre-European state. The Boolgooroo wetland covers an area of 60 ha of the total 160 ha Mungalla wetland complex (Fig 1). The wetland is bounded to the west by grazing lands and to the east by regrowth forest on coastal sand ridges. Mangroves and saltmarsh patches are present along the coast and to the south of the property, which is adjacent to the Great Barrier Reef lagoon. Inland, the surrounding catchment is dominated by sugar cane farms, with some areas of grazing.

The area has a wet tropical climate with highly variable seasonal and annual rainfall. The long term mean rainfall at nearby Allingham is 2060 mm and is strongly seasonal with 85% falling in the six wettest months, November to April. Temperatures are highest in December (daily average 29.1°C) and lowest in July (daily average 20.4°C), with high humidity (~63 – 77%) throughout the year.

Because of the highly seasonal rainfall, freshwater only enters the wetland in the wet season as direct rainfall input, runoff from the surrounding sub-catchments and overbank flow from Palm Creek, which runs along the western boundary of Mungalla station. Once the bund was removed, the upstream wetland was again connected to the coastline, with the possibility of being fed tidally with seawater from a tributary of Palm Creek (just south of Forrest Beach) and Cassady Creek (Fig 1).

### Vegetation monitoring

Vegetation monitoring and mapping in the wetland was carried out using Worldview-3 8-band satellite imagery (DigitalGlobe Inc. Longmont, CO, USA) pan sharpened to a resolution of 0.31m. Imagery was collected once each year between August and September over the Mungalla station region, then cropped to an extent around the Boolgooroo wetland (Fig 1). Classification of the imagery into major vegetation types was carried out using object-based image classification techniques [46], using nearest neighbour supervised classification, and manual classification in more difficult areas. Complexity of the wetland area required separate classification of small regions, incorporating different segmentation levels within the object-based process, which were later merged. To enable the supervised and manual classification, dominant species were recorded at 50 fixed ground truthing sites across the imagery extent each year. In addition, high-resolution aerial photography was obtained with a Go-Pro camera attached to the underside of a helicopter flying along several transects that ran parallel and across the wetland site from west to east and at an elevation of approximately 100 m. These images were geo-referenced to preselected ground control points (geo-located with a differential GPS). Major vegetation groups could be clearly identified visually from the high-resolution aerial photography which was used as further ground truth in classification of the Worldview-3 imagery. A comparison of final classification imagery was carried out between consecutive collection dates (Aug/Sep yearly), to identify differences in the area inhabited by these major vegetation groups.

### Water depth and quality measurements

Wetland water depth, temperature and electrical conductivity were monitored by loggers (CTD-Diver, Eijkelkamp Soil & Water, The Netherlands) located in five permanent positions in the wetland, beginning on 24^th^ October 2012. The locations are 450 m, 250 m and 50 m above, and also 50 m and 250 m below the bund wall (Fig 1). The above bund logger locations were all within the weed infested parts of the wetland. The loggers captured data from the bottom of the water column (10 cm above the soil surface) every 15 minutes and were downloaded as part of monthly routine maintenance visits. An additional water quality logger (Hydrolab, OTT Hydromet, Colorado, USA) was used above the bund at the 50 m location, adjacent to the Diver logger (Fig 1). The Hydrolab logger recorded values of electrical conductivity, depth, pH and dissolved oxygen concentration every 30 minutes at 10 cm above the soil surface. Ancillary data used in conjunction with the above wetland monitoring data are daily rainfall measured at Allingham (Australian Bureau of Meteorology (BOM) station No 032117) and tide data recorded at the Lucinda Jetty; which was then interpolated to Forrest Beach; 1.5 km east of the wetland (4min offset). The tidal delay from the coast at Forrest Beach to the bund position is approximately 3 hours. Further details of the tidal interpolation method are given by Karim et al, [47].

### Fish biodiversity

Prior to this project a fish survey was conducted in the wetlands using baited collapsible box traps along with cast nets, electrofishing, and visual observation. Sampling occurred at sites among the invasive vegetation [48]. For comparison we conducted another fish survey with the use of a boat mounted electrofisher (Smith-Root 2.5 GPP generator mounted vessel) in May 2016. In this second survey, approximately one third of the wetland was surveyed over a 55 min period, around fringing vegetation and woody debris (the remaining two thirds of the wetland were too shallow for the boat to access). All native fish were identified and immediately released after capture, while pest fish species were euthanised using accepted methods. Given the differences in the sampling methods before and after bund wall removal, the data presented represents a presence/absence of the catch records.

## Results

### Before Bund removal

#### Vegetation monitoring

Prior to bund removal (2012/13), with no changes in EC above the bund, over two thirds of the area were inhabited with ponded pastures and exotic floating vegetation. A small percentage was inhabited by *Nymphaea gigantea:* a native water lily during 2012 which increased into open waters during 2013 (Table 1) (Fig 2(a,b)). Over half of the area was infested with weeds of national significance (WONS) (Table 1), dominated by *Hymenachne amplexicaulis* (Olive Hymenachne) and *Eichhornia crassipes (Water Hyacinth).* On average open water made up less than one fifth of the Boolgooroo area (Table 1).

**Fig 2.**
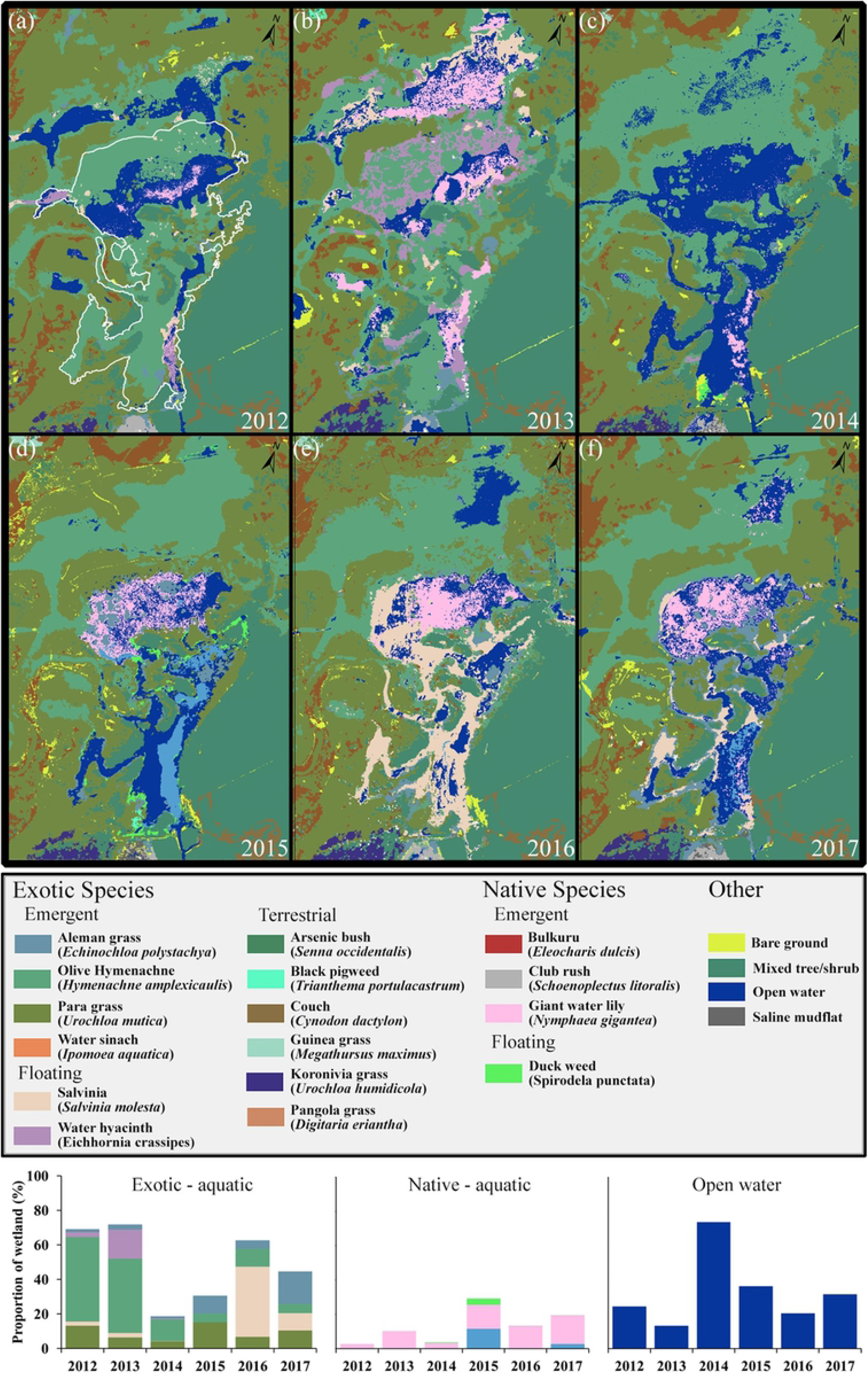
Changes in vegetation from 2012 to 2017. The white outline in (a) represents the Boolgooroo region on which change analysis was based. Charts on the bottom of the figure represent changes in exotic/native vegetation and open water (colours correspond to the classification key).

**Table 1.** Area inhabited by vegetation groups (%): pre- (2012/13) and post (2014 – 2017) bund removal.

| Satellite Classification<br>(% of Boolgooroo site) | Survey Year |  |  |  |  |  |
| --- | --- | --- | --- | --- | --- | --- |
|  | 2012 | 2013 | 2014 | 2015 | 2016 | 2017 |
| **Exotic aquatic plants** |  |  |  |  |  |  |
| Aleman grass<sup>†</sup><br>(*Echinochloa polystachya*) | 1.85 | 3.12 | 1.18 | 10.26 | 4.87 | 18.93 |
| Olive hymenachne*<sup>†</sup><br>(*Hymenachne amplexicaulis*) | 48.94 | 43.04 | 12.61 | 5.11 | 10.51 | 5.23 |
| Para grass<sup>†</sup><br>(*Urochloa mutica*) | 13.26 | 6.45 | 4.24 | 15.16 | 6.83 | 10.42 |
| Salvinia*<br>(*Salvinia molesta*) | 2.41 | 2.61 | 0.04 | 0.00 | 40.53 | 10.12 |
| Water hyacinth*<br>(*Eichhornia crassipes*) | 2.68 | 16.72 | 0.55 | 0.03 | 0.00 | 0.00 |
| **Native aquatic plants** |  |  |  |  |  |  |
| Bulkuru<br>(*Eleocharis dulcis*) | 0.00 | 0.00 | 0.00 | 11.46 | 0.26 | 2.59 |
| Club rush<br>(*Schoenoplectus litoralis*) | 0.00 | 0.00 | 0.00 | 0.00 | 0.00 | 0.10 |
| Duck weed<br>(*Spirodela punctata*) | 0.00 | 0.00 | 0.31 | 3.57 | 0.00 | 0.00 |
| Water lily<br>(*Nymphaea gigantea*) | 2.66 | 10.16 | 3.31 | 14.07 | 12.91 | 17.54 |
| **Other** |  |  |  |  |  |  |
| Open Water | 24.56 | 13.38 | 73.71 | 36.38 | 20.56 | 31.70 |

\*Weeds of national significance, <sup>†</sup>Ponded pasture grasses.

#### Water depth and quality measurements

Towards the end of 2012 wetland depth at all locations declined steadily with only 0 to 20 cm of water depth above the bund and even less (0 to 10 cm) below the bund (Fig 3a,b) by the middle of December 2012. Subsequent rainfall (Fig 3c) increased the wetland depth, which reached a maximum of ~170 cm (above and below the bund) at the end of January 2013. After February 2013, water levels dropped rapidly, and then more slowly as drainage from the wetland slowed. Smaller rainfall inputs to the wetland in March, April and May 2013 resulted in only modest increases in water depth (~ 10 cm). After this time water depth again dropped steadily, approaching zero in October 2013 just before the bund was removed. There were multiple occasions throughout the 2013 dry season when depth increased, at approximately monthly intervals, at 250 m and 50 m below the bund (Fig 3b). However, there was no corresponding increase in depth above the bund on any of these events (Fig 3a); it would therefore appear that the depth increases below the bund were due to tidal pulses and that these did not penetrate above the bund at this time.

**Fig 3.**
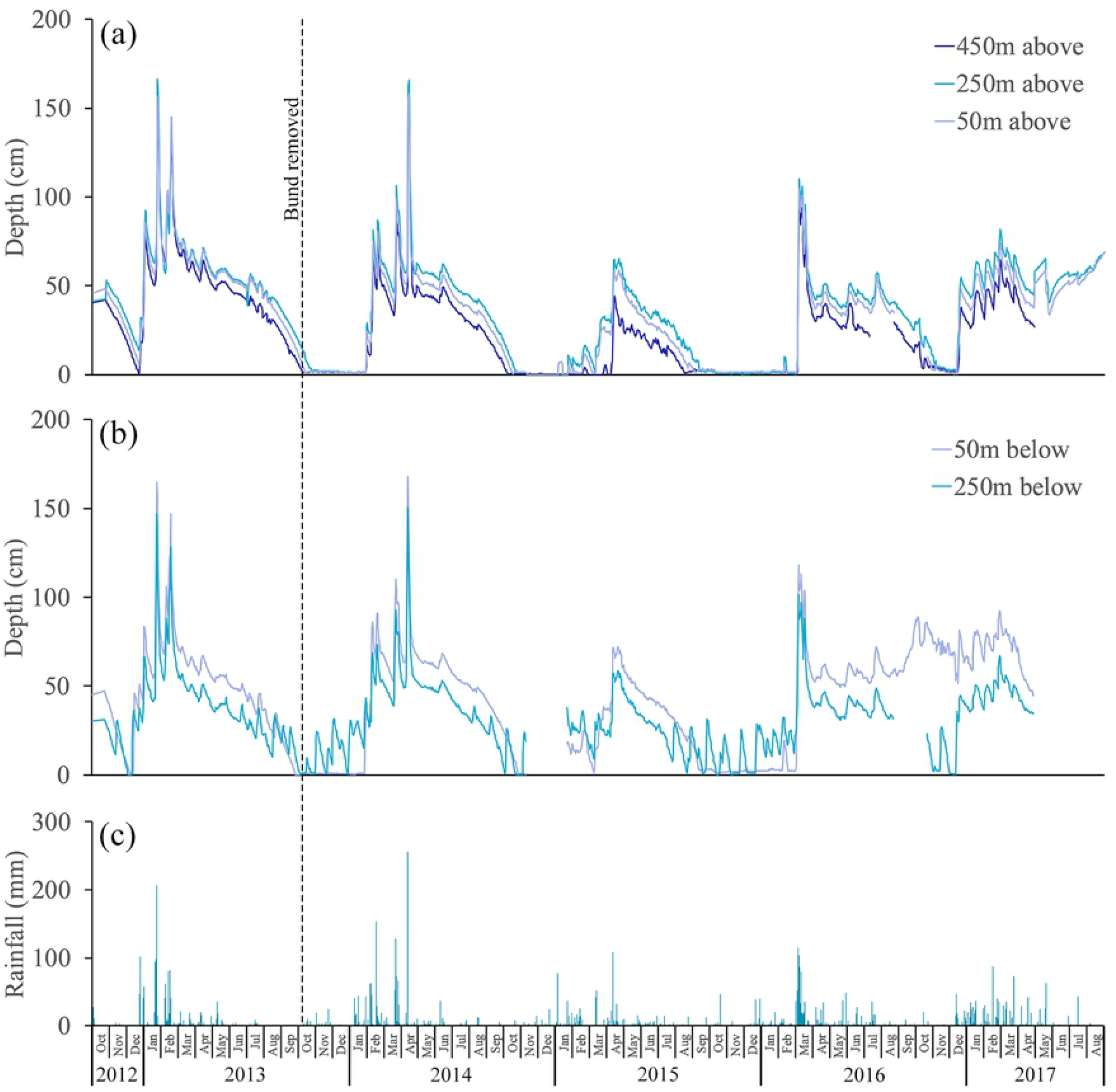
Changes in depth above and below the bund location and daily rainfall – October 2012 to August 2017. Logger distances above the bund (a) are 450 m, 250 m and 50 m and below (b) are 50 m and 250 m. The bund was removed on the 6^th^ October 2013. Daily rainfall (c) was derived from the nearby Bureau of Meteorology station at Allingham approximately 2.5 km distant.

Seasonal variations in wetland electrical conductivity (EC) are shown in Fig 4a,b. Before removal EC above the bund was never greater than 0.5 mS cm^−1^ (Fig 4a). During the same period there were 9 events where EC increased below the bund (Fig 4b), sometimes approaching seawater (~ 55 mS cm^−1^). Each event consisted of >=2 consecutive days of high tides (Fig 4c). Tidal water reached the below 250 m location more often (9 times) compared to the below 50 m location (2 times), demonstrating that the gentle slope between the two locations is sufficient to reduce the frequency and duration of tidal water approaching the bund. The smallest tide to reach the below 50 m location was ~3.7 m, indicating that tides needed to be equal to or greater than this to reach the bund location.

**Fig 4.**
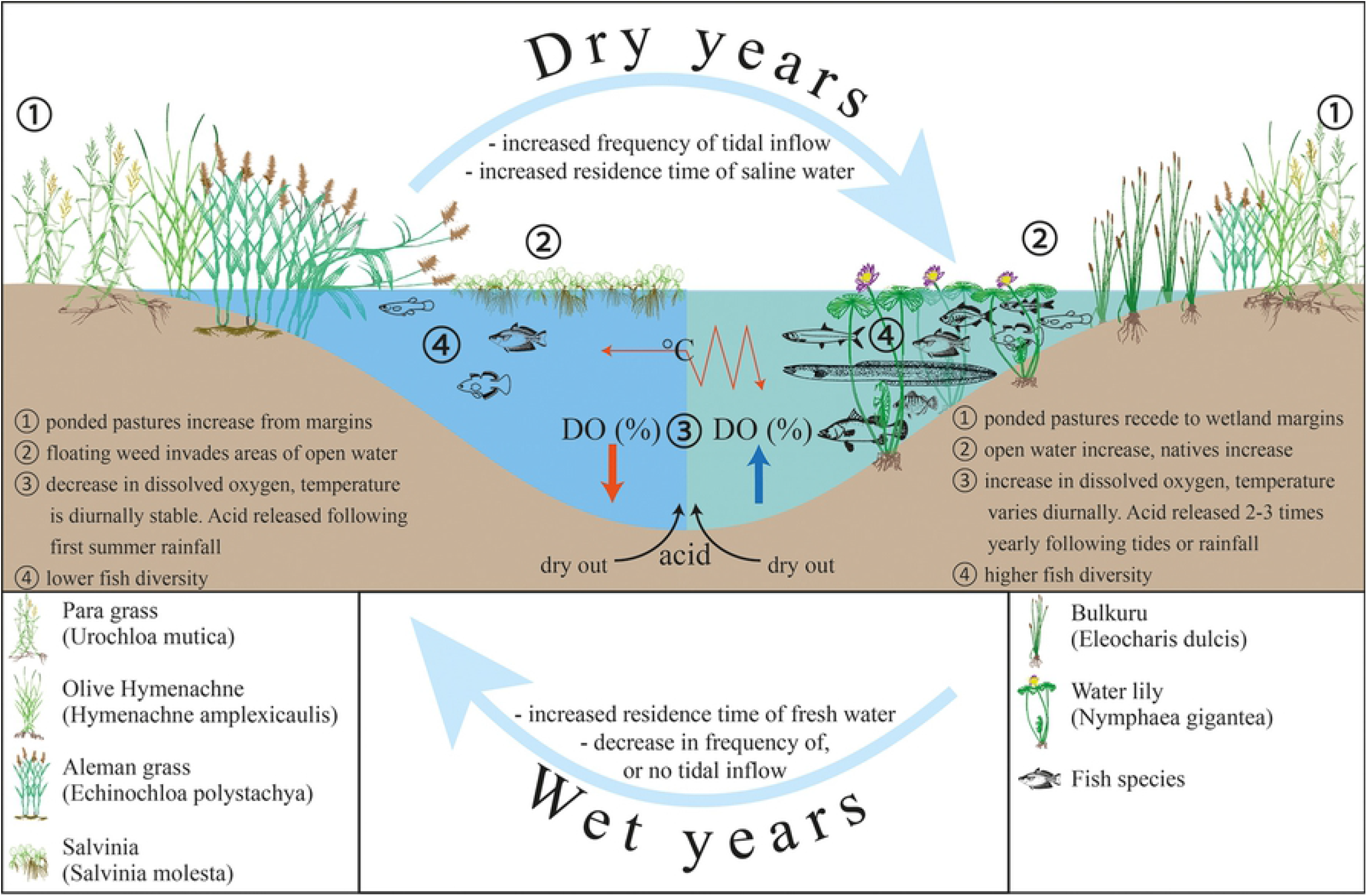
Changes in salinity above and below the bund location and tides >=3.7 m – October 2012 to August 2017. Logger distances above the bund are 450 m, 250 m and 50 m and below are 50 m and 250 m. The bund was removed on the 6th October 2013. High tides (>= 3.7 m) are for Forrest Beach.

Dissolved oxygen (DO) (recorded 10 cm above the bottom of the wetland) was consistently close to zero before the bund was removed and pH showed little variation prior to bund removal being slightly acidic averaging ~6 (Fig 5a,b). Temperature showed little diurnal oscillation at this time (Fig 5b).

**Fig 5.**
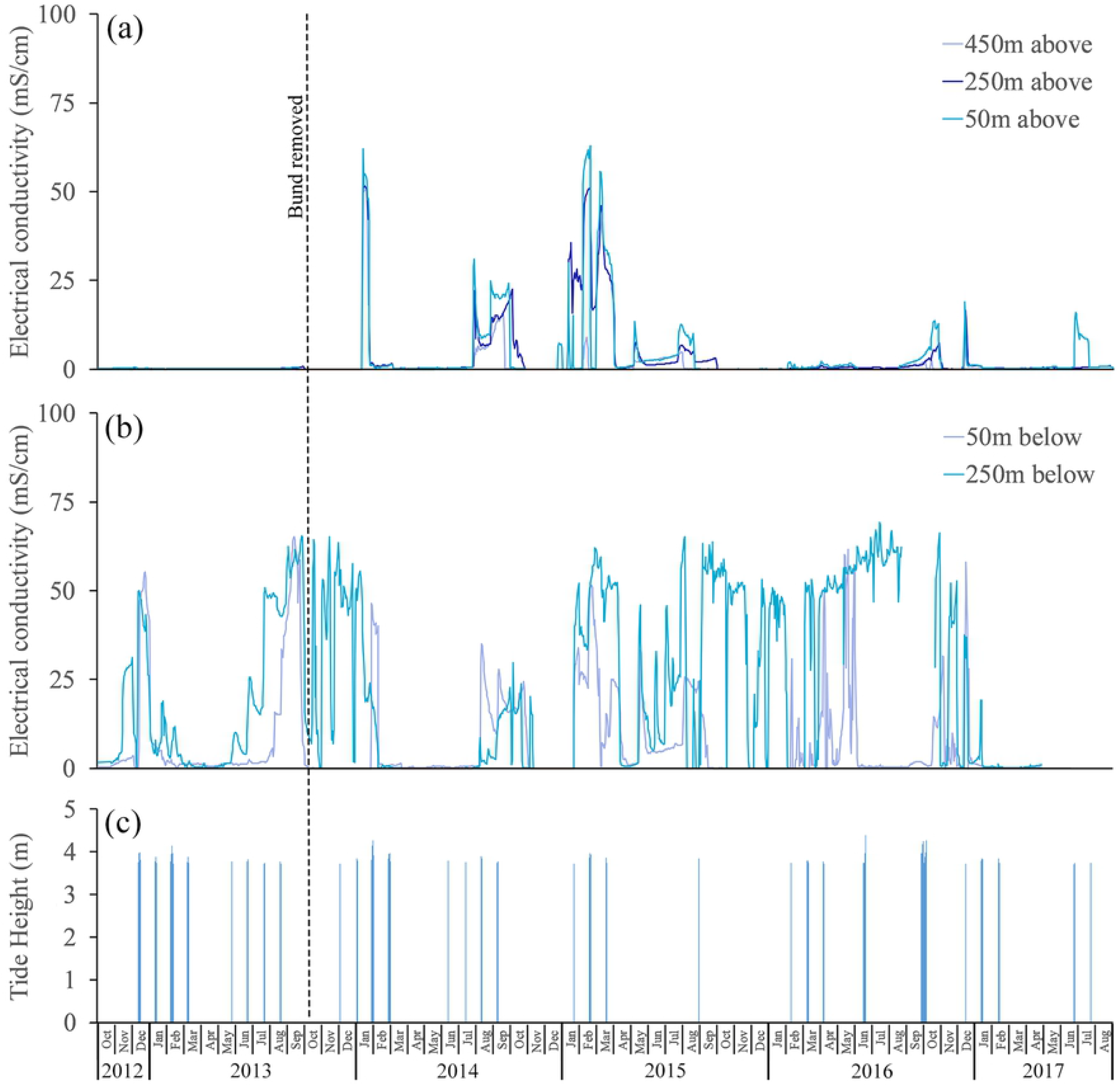
Seasonal changes in dissolved oxygen, pH and temperature. Daily average dissolved oxygen (% saturation) and pH were recorded by the Hydrolab logger at the above 50 m location (a). Daily average water temperatures shown are as measured 250 m above the bund location (b). Diurnal oscillations in dissolved oxygen (30-minute intervals) and water temperature (15-minute intervals) are also shown (a), (b).

#### Fish biodiversity

The fish community in Mungalla wetland prior to bund wall removal was of low abundance and diversity in comparison with other coastal freshwater wetlands in the region [49], with only three species recorded (Empire gudgeon, *Hypseleotris compressa;* Eastern rainbow fish, *Melanotaenia splendida inornata;* the invasive Mosquitofish, *Gambusia holbrooki)* despite sampling efforts comprising a combination of cast nets, bait traps, electrofishing and visual observation [48] (Table 2). The most abundant species was the empire gudgeon *(Hypseleotris compressa)*, which is widespread in the region owing to a diadromous movement ecology where it migrates to access upstream and downstream river areas [50]. The high abundance of the empire gudgeon suggests either it is capable of circumventing the earth bund wall, and/or it can tolerant poor water quality, as has been the case in the wetlands prior to bund wall removal.

**Table 1:** Fish community composition in the Mungalla wetland (Boolgooroo) before bund removal in 2009 and following removal in 2016.

| Common name<br>Species | Pre bund removal survey<br>(2009) | Post bund removal survey<br>(2016) |
| --- | --- | --- |
| Mosquitofish<br>*Gambusia holbrooki* | ✓ | ✓ |
| Hardyhead<br>*Craterocephalus stercusmuscarum* |  | ✓ |
| Empire gudgeon*<br>*Hypseleotris compressa* | ✓ | ✓ |
| Tarpon<br>*Megalops cyprinoides* |  | ✓ |
| Rainbowfish<br>*Melanotaenia splendida* | ✓ | ✓ |
| Barramundi*<br>*Lates calcarifer* |  | ✓ |
| Silverbiddy*<br>*Gerres subfasciatus* |  | ✓ |
| Banded scat*<br>*Selenotoca multifasciata* |  | ✓ |
| Long finned eel*<br>*Anguilla reinhardtii* |  | ✓ |
| **Total** | 3 | 9 |

\*diadromous movement ecology

### After Bund removal

#### Water conductivity and effects on vegetation

Following bund removal (6^th^ October 2013), re-establishment of connectivity to Cassady creek was immediately evident with slow freshwater outflow over 12 days to reach an equilibrium depth of ~10 cm lower than the bunded state. Water and vegetation condition of the wetland remained unchanged for the following 3 months.

In January 2014 the wetland experienced the first tides over 3.7 m (3.8 and 3.7 m) which did not affect the wetland above the bund location, these were followed by 5 tides over 3.7 at the end of the month (Fig 4c). These tides were unusually high due to a severe tropical low in the Coral Sea at that time. As a result, the wetland became influenced by tidal ingress (> 46 mS cm^−1^) and remained so for the next 10 days before rainfall again suppressed conditions (Fig 3a,c). Conductivity levels remained brackish (> 1 < 46 mS cm^−1^) for the next 7 days before returning to freshwater (< 1 mS cm^−1^). In 2014 there were 7 tidal periods (most with 3 consecutive tides over 3.7m) that affected the wetland. The wetland retained tidal water signal for an average of 10 days, becoming brackish for an average of 118 days. Three of these tidal periods were recorded at the 450 m above bund location.

The effect of increased EC on the ponded grasses and weeds was noticeable from a few days after the first tidal ingress at the end of January 2014, with a rapid yellowing of the vegetation occurring. Weed death continued after this and by the time of the next remote vegetation survey in August 2014 ponded grasses and other exotic species had been greatly reduced to below 20% within the tidally effected area (Table 1). *Nymphaea gigantea* also decreased by two thirds at this point (Fig 2c; Table 1). A dramatic change is evident (Fig 2c) with most of the exotic vegetation being replaced by open water.

The following year (2015) saw 4 tidal periods over 3.7 m with the wetland becoming more saline for an average of 13 days and then brackish for 178 days, again with 3 periods being recorded at the 450 m above bund location. 2015 was the driest year on record for the region, which accounts for the differences in tidal inundation (days), as wetland depth and associated freshwater outflow did not restrict tidal inflow to the same degree as the previous year.

Satellite imagery recorded an increase in two of the ponded pastures – Aleman (E. *polystachya)* and Para grass *(U. Mutica)* – invading areas previously inhabited by Olive Hymenachne *(H. amplexicaulis)* this occurred on the wetland margins and comprised ~25% of the site (Fig 2d, Table 1). Olive Hymenachne was reduced to ~5% with Salvinia and Water Hyacinth being undetectable on the remote sensing imagery (Fig 2d) (Table 1). There was an increase in native aquatic plants driven by the appearance of *Eleocharis dulcis* (Bulkuru) within the lower half of the site as well as an increase in *N. gigantea* in the upper site. *Spirodela punctata* (Duck weed) was also present (~3.5%) in some of the shallower areas originally inhabited by *H. amplexicaulis.* The native species generally took advantage of the disappearance of exotic species and newly available open water.

Water quality showed improvements post weed removal (Fig 5a). pH (6.5 – 8.4) and DO (60 – 120 % saturation) were most frequently maintained within ranges expected in wetlands in the region [49]. The improvements in DO seen here are similar to those found by Perna and Burrows [51] following mechanical removal or water hyacinth mats in the lower Burdekin region of north Queensland. There were still short periods where oxygen fell below acute and chronic thresholds for biota [52] however, none were as low or lasted as long as seen prior to bund removal. Two occasions were recorded in early 2015 when pH became acidic (~3.4) which could adversely affect survival of aquatic biota, the most likely cause was leaching of sulphuric acid following oxidation of acid sulphate soils after drying out in the upper Boolgooroo at the end of 2014. These levels were short lived and pH was restored to an average of ~7.4 after buffering by further tidal intrusion and rainfall.

The entire wetland dried out in October 2015 remaining dry for two months (Fig 3a) and then received small flushes of rainfall until March 2016 (pH as low as ~3 again occurred due to acid sulphate soil oxidation and leaching). At this time, as a result of increased wetland accessibility, predation by feral pigs removed most of the Bulkuru *(E. dulcis)*, a common and damaging event in northern Australia [53–55].

Although there was a single high tide of ~3.7 m in February of 2016 it did not enter the wetland (Figure 4a,c). Similarly, this also occurred in early January of 2014 indicating that not only is ~3.7 m is the lower limit for tidal flow into the wetland it will only occur when there are 2-3 consecutive tides above this level or when freshwater depth and outflow is very low as in January of 2015. There was a large amount of rainfall during March of 2016 with constant rainfall input until July. The wetland remained as freshwater until September/December when tidal water reached upstream areas of the bund wall, resulting in small conductivity peaks. 2016 also saw the return of *S. molesta* (~40% of the area) most likely due to the prolonged period without tidal input, invading open water and the areas previously inhabited by *E. dulcis and N. gigantea* in the upper reaches *(Fig 2e). E. polystachya* and *U. mutica* were reduced slightly from the margins to be replaced by *H. amplexicaulis.* Whilst some of the area occupied by *N. gigantea* were now used by Salvinia, the water lily had become denser (Fig 2e) and remained stable at around ~13% of the Boolgooroo area (Table 1). *E. dulcis* was no longer detectable on the imagery after the feral pig predation event of 2016, however some individual plants were observed in the lower Boolgooroo area at this time. pH dropped to ~3 during the rainfall period in March following on from the dry out that happened in late 2105 – indicating again that the lowering of pH is a product of acid sulphate soil oxidation and sulphuric acid release on wetting, it then maintained an average of around 6.5 for the remainder of the year (Figure 5a). Available DO dropped to close to zero after the March 2016 rainfall, most likely due to the rapid increase of Salvinia – growth of which may have been accelerated due to the low pH levels [56]. Mats of Salvinia typically lower oxygen levels by inhibiting oxygen transfer across the water surface. This occurs in particularly slow flowing situations such as those seen in the Mungalla wetland [57–59].

In the final 7 months of the study (2017) 3 tides affected the wetland, but only to the above 50 m location (Figure 4a) although the spikes in EC from September and December 2016 tides could be considered here in regards to vegetation change as tidal input during this period was observed to drastically reduced the population of Salvinia, which is particularly sensitive to even low levels of salinity [60, 61]. The 2017 image classification (Fig 2f) clearly shows the reduction in Salvinia to marginal locations with *N. gigantea* moving into the areas that it previously occupied in the upper Boolgooroo, and Bulkuru reappearing in the lower Boolgooroo. At this time the saltmarsh species *S. litoralis* (Club rush) appeared in a small area (Table 1) directly above eastern end of the bund location (Fig 2f). Ponded grasses, which had remained on the wetland margins since 2014, showed a decrease in *H. amplexicaulis* down to ~5%, similar to the 2015 level and may represent the effective control extent of tidal intrusion. Growth of *U. mutica* and *E. polystachya* increased to inhabit areas previously occupied by *H. amplexicaulis* in 2016, with the increase by Aleman grass (E. *polystachya)* even more pronounced in this period. Typical of the coastal seawater, pH rose to a level just below 9 following the tidal inflow of September 2016 to February 2017, it then gradually fell to ~7.5 in May of 2017, dissolved oxygen reached a maximum level of ~100 % saturation during the period between November 2016 and May 2017 coinciding with rainfall events, however, the average DO was ~2 % saturation with many readings close to or at zero. Unfortunately, both loggers below the bund location stopped recording correctly around February of 2017, and the Hydrolab failed in May of 2017 (Fig 3b,4b,5a) – data collection from those loggers was discontinued at those points.

#### Dissolved oxygen and water temperature

Dissolved oxygen saturation (recorded 10 cm above the bottom of the wetland) was consistently close to zero before the bund was removed (Fig 5a). However, in the first wet season after its removal, DO improved following the first series of seawater pulses that entered the wetland in January and February 2014 (Fig 4c), reaching ~ 100% saturation on most days (Fig 5a). DO declined again for several weeks after this until the freshwater pulses in March and April 2014 (Fig 3a) again improved DO; however, initially there were relatively few days when DO reached 100% saturation. By June and July 2014 DO improved further, approaching 100% saturation on many days. As wetland depth dropped below 50 cm in August and September 2014, the seawater pulses that could now enter the wetland continued to sustain reasonably high DO, and there was even a supersaturated spike (~ 200%) in DO about two weeks after the seawater ingress in August 2014. Despite the low water levels in January and February 2015, DO concentrations reached 80 to 100% on most days, however much lower DO was recorded during March, April and May 2015.

Water temperature in the wetland varied seasonally, being coolest in June/July (~ 22.5°C) and warmest in January/February (~ 29.5°C) (Fig 5b). Water temperatures at the three locations above the bund were very similar, but the water was slightly warmer (by ~ 1°C) below it. Daily average water temperatures in 2014 to 2017 were similar to those recorded in 2012/2013 before bund removal. The highest temperatures were recorded in January/February 2015, averaging ~31°C above the bund location, compared to ~29°C and ~29.5°C in the two previous summer seasons, and by ~30 °C and ~29 °C in the following two seasons. The higher temperatures in 2015 coincided with the low rainfall received that summer season, leaving the wetland very shallow, averaging only 11 cm above the bund location in January and February (Fig 3a).

Water temperature also varied diurnally, generally by 1 to 3 degrees, but there were also notable periods when the diurnal oscillations increased to > 10 degrees; for example, during the period October 2013 to January 2014 and for most of the 2014-16 seasons and again during the summer months in 2016/17 (Fig 5b). Large diurnal oscillations in temperature coincided with periods of shallow wetland depths. The size of the diurnal oscillation is important as it defines how long aquatic species in the wetland are exposed to the highest water temperatures during the afternoon.

The thermal regimes 250 m above and 250 m below the bund are compared in (Fig 6). These plots show how often water temperature exceeded a given temperature threshold and are compiled from all 15-minute recordings made from the warmest time of the year in January, February and March (JFM) 2013 (before bund removal) and the same three months in 2014 and the driest year 2015 (after bund removal).

**Fig 6.**
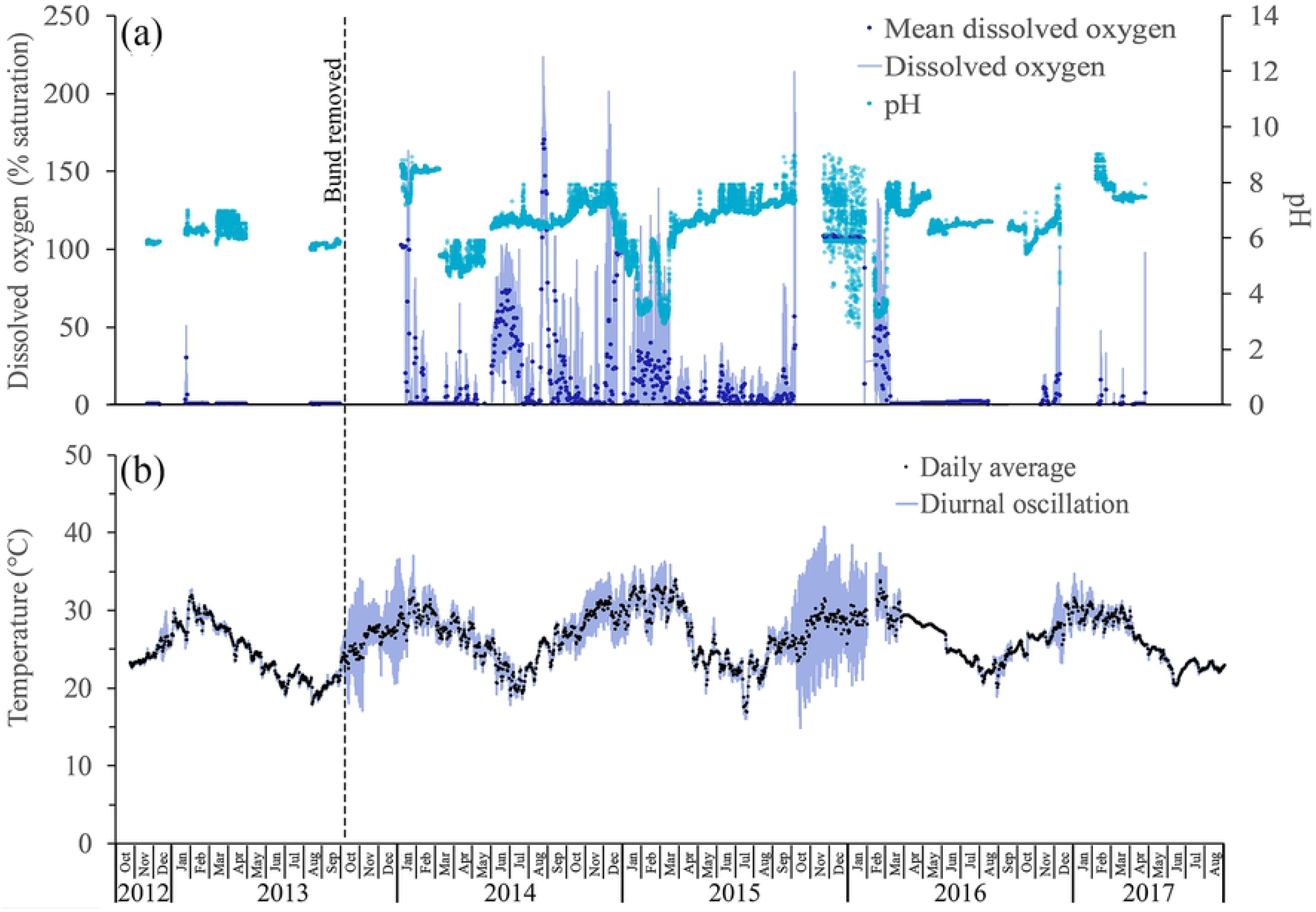
The percentage of time water temperature exceeded any given temperature threshold during the warmest months (January, February and March) in the Mungalla wetland (Boolgooroo). (a) compares the temperatures 250 m above (black) and 250 m below (grey) the bund before it was removed. (b) compares the temperatures 250 m above the bund before (black) and after (2014; short-dashed black; 2015; long-dashed black) it was removed. The exceedance of threshold T_pref_ = 31°C is also shown

Water above the bund was above 26°C during JFM 2013, but rarely exceeded 32°C (Fig 6a). Water was warmer below the bund during this period, reaching 35°C. The exceedance of the preferred temperature threshold for some tropical freshwater fish (see [62]), *T_pref_* = 31°C, was much higher below the bund (23% of the time) compared with above the bund (6.9% of the time). Comparison of 2013 and 2014 thermal frequency curves (Fig 6b) shows that bund removal *per se* did not markedly affect the wetland thermal regime. However, water temperature was very dependent on (shallow) wetland depths, which were mainly determined by the timing and amount of wet season rainfall. This was the situation in the 2014/15 season where the low rainfall led to shallow wetland depths (Fig 3a). As a result, diurnal temperature oscillations were very high (Fig 5b) and led to prolonged periods of high temperature, as shown by the temperature frequency curve for JFM 2015 (Fig 6b). For example, the exceedance of the temperature threshold *T_pref_* = 31°C, was 61% in JFM 2015, compared with 7.3% and 6.9% in 2014 and 2013 respectively. Temperatures never exceeded 34°C in 2013 and 2104, but this temperature was exceeded 34% of the time in 2015, and even approached 40°C at times.

#### Fish Biodiversity

Post bund removal fish surveys conducted in May 2016, found the same three species identified before bund removal with an additional six species recorded. While species count is still low by local standards [49], the increase in species number has presumably been the result of reconnecting the wetlands with the downstream estuary. This along with the removal of aquatic weeds and associated improvements in pH and dissolved oxygen has allowed more species to access and live in the wetland area. Increases in fish community richness have been recorded elsewhere in the region following aquatic weed removal [35]. The most notable records in the post bund survey were barramundi *(Lates calcarifer)*, silver biddy *(Gerres subfasciatus)* and banded scat *(Selenotoca multifasciata);* each of these species have a diadromous movement ecology, requiring access to wetlands to fulfil critical life cycle stages.

## Discussion

The restriction of seawater from coastal wetlands using earth bunds (e.g. dams, dykes or levees) is widespread and has been seen to cause the degradation and/or loss of salt marsh ecosystems worldwide [3, 9, 63]. Prolonged exclusion of seawater leads to the loss of native halophytes and widespread invasions of freshwater species, in addition to major changes in sedimentation rates and soil chemistry [64]. Recognition that these bunded wetlands are not natural has led to an increasing number of tidal restoration projects [9]. For example, Smith *et al.*, [65] describe how freshwater plants are being replaced by native salt marsh plants following the restoration of tidal flows to the Hatches Harbor salt marsh in Cape Cod, Massachusetts, USA. In Australia, tidal flows were reinstated into coastal wetlands in the Hunter estuary in New South Wales and this increased the area of salt marsh, largely through expansion into areas of pasture [27]. Conversely seawater intrusion into coastal wetlands such as the Gippsland lakes in Victoria [66] and the rivers in the Northern Territory [67] has also caused changes in wetland ecology, however, these are generally seen as undesirable, since in these cases they changed the wetlands from their natural freshwater state. These salinization effects are reported worldwide and are attributed to a range of anthropogenic impacts including sea level rise [68]. In the case of the Boolgooroo wetland on Mungalla station the removal of the earth bund and subsequent reintroduction of tidal water ingress, returning the wetland to its historical halophytic state was a desirable outcome, where it would receive occasional tidal pulses, enough to assist with naturally supressing invasive aquatic freshwater plant species.

Seawater entry into the Mungalla wetlands does not occur often since, with the bund removed, only the highest of tides, of approximately 3.7 m, are able to penetrate the wetland. These usually occur in sequences of 2 to 4 consecutive days on around four occasions during the summer (December to March) and a similar number in winter (June to September). However, if the wetland contains water deeper than ~ 0.5 m a ‘hydraulic barrier’ is formed meaning that tides effectively need to be greater than 3.7 m to penetrate the wetland upstream of the bund wall location. This may not happen frequently as tides in excess of 4.0 m only occur about once a year. Seawater ingress is therefore more likely to happen when the wetland depths are lowest, either during winter or in years of low summer rainfall (which were experienced during the research program here). However, when hydraulic pressure is low during low rainfall years and in the pre-summer period the frequency and duration of seawater ingress to the Boolgooroo region of the Mungalla wetland can exceed that required to cause permanent damage to the invasive freshwater weeds. Salinity tolerance tests carried out by Reid et al., [69] found that the growth and survival of Aleman grass *(Echinochloa polystachya)*, Olive Hymenachne *(Hymenachne amplexicaulis)*, Para-grass *(Urochloa mutica)* and Water hyacinth *(Eichhornia crassipes)* were all affected by changes in EC, even when exposed to only 30% seawater concentration for as little as a single day. Although tolerance varied between these four weed species (Aleman grass being most tolerant), each had stopped photosynthesis and mortality rates were very high when exposed to 100% seawater equivalent for longer than 7 days. Clearly removal of the earth bund allowed seawater to penetrate well into the wetland on multiple occasions creating saline conditions that would be expected to have a marked impact on the freshwater adapted weeds found above the bund location. Indeed, surveys of the wetland vegetation before and after the removal of the bund show that there was a large and relatively rapid reduction in freshwater weeds above the bund location, with the re-emergence of native salt tolerant plants after only 2-3 years. However, since wetland depth and tidal height conditions required to change the wetland vegetation towards a more halophytic composition may only occur in the driest of years (as in 2015), or during dry pre-summer conditions, there is a risk of the wetland reverting to dominance by invasive freshwater species. In fact, this occurred in 2016 with an initial reinvasion of the floating species *S. molesta* (Salvinia) followed by some encroachment of ponded pastures from the wetland margins. Of great interest, following saltwater ingress, has been the expansion of Aleman grass (E. *polystachya)*, being the most tolerant to saline conditions [69], which will most likely continue to dominate from the wetland margins inwards, replacing Para grass *(U. mutica)* and Hymenachne *(H. amplexicaulis)* when conditions are suitable.

Fish biodiversity increased after bund removal, occurring as a result of amended connectivity between the wetland and the ocean and improved water quality conditions. The very poor water quality in the wetland, especially the extremely low dissolved oxygen which would regularly expose, such as fish (and other aquatic fauna), to acute and chronic hypoxia risks led to low fish numbers in the wetland prior to bund removal. The low oxygen levels before bund removal were likely to have been due to the presence of the dense mat of freshwater weeds that limited oxygen transfer and light penetration to the bottom of the wetland. Butler and Burrows [52] found that there were significant risks of acute exposure when DO was below 30% saturation (the ‘acute trigger value’ ATV). Prior to bund removal hypoxia risks were severe, as DO was below the ATV threshold virtually 100% of the time. With the earth bund removed, DO gradually increased in the first wet season (2013/14), but still fell below the ATV for 93% of the time. The improvement in DO was more dramatic following the 2014 dry season, where DO only fell below the ATV for 49% of the time. Given the normal diurnal cycling of DO at the bottom of a wetland [70], these latter trigger value failures should not present a high risk to fish that can swim towards the surface where DO would be higher [71]. An aquatic ecology survey of the Mungalla wetlands completed 4 years before the bund was removed [71] found that the weed infested sections of the wetland did not provide suitable habitat refugia for most fish species, and created conditions that increase the wetlands’ susceptibility to acute episodic periods of low dissolved oxygen. That study also showed that water quality was better in ‘open’ (weed free) parts of the wetland. This was probably the habitat (22% of the wetland area) where the few species of fish (3 species) existed at that time (May 2009). Reduction of wetland depth following bund removal may also pose a risk to wetland biota via its effect on water temperature. Water temperatures recorded above the Mungalla bund location did not exceed 34°C either before (2013) or after (2014) removal. Therefore, bund removal *per se* did not affect the wetland thermal regime. However, when water depths were very shallow (< 40 cm), as in the 2015 wet season, due to the very low rainfall in that year, temperatures exceeded 31°C over 60% of the time, 34°C for 35% of the time. These higher temperatures could have had a major impact on freshwater aquatic species, if trapped within the wetland, as many tropical fish [72] and freshwater crustaceans [73, 74] cannot survive prolonged exposure to such high temperatures. However, with connectivity improved via bund removal and a reduction of weed species, aquatic biota could move to other parts of the wetland or adjacent creeks during these periods. This is also true for the short periods of low pH when the wetland water became acidic (pH 3 – 3.5), reaching conditions that could be lethal for aquatic invertebrates and could even kill fish [75, 76]. These conditions follow periods of drought, when the acid sulphate soil in the wetland is exposed to air, releasing a pulse of sulphuric acid [77] when the wetland is initially reflooded by fresh or saline waters. Interestingly, if high tides continue to occur during these periods of low wetland depth, the alkaline seawater ingress can eliminate the acidic conditions; which would not have occurred when the wetland was bunded. However, the buffering of saltwater potential has been shown to contribute to secondary implications, including the precipitation of heavy metals that are available in the water column [8].

Freshwater weed infestation is widespread in many of the coastal wetlands in North Queensland [32, 35], the primary control mechanisms are aerial spraying with herbicide and mechanical removal [35, 78]. However, these methods are expensive, can have undesirable ecological consequences and only effective for a limited time. In some cases, these methods have been shown to have little to no impact (Hurst and Boon, 2016). For example, in the years before bund removal on Boolgooroo chemical spraying with herbicide did increase the open water area, but this mitigation measure provided only a temporary solution with aquatic weeds such as Olive hymenachne again present only 2-3 months after spraying and Aleman grass largely unaffected most likely due to herbicide resistance at the application rates used [79]. Although spraying with more ecologically acceptable saline water has been attempted for control of Water hyacinth [80] it is not yet widely used for freshwater weed control. With approximately 30% of coastal wetlands being bunded in the Great Barrier Reef region [81], removal of an earth bund or levee could provide a more cost effective and sustainable means of controlling freshwater weeds and improving water quality. However, landholders and government do still need to take care to fully consider tidal boundary laws and amendments when considering ponded pasture reconversion projects [82].

## Conclusions

With the limited success of control methods to restore wetlands as productive coastal features in the Great Barrier Reef catchment area, this study revealed that reinstatement of tidal flows into bunded estuarine wetlands is relatively effective in the removal of freshwater weeds and ponded pastures. As a passive remediation method reintroduction of tidal flow is a sustainable, efficient and cost-effective management option for restoring aesthetic and ecological values of coastal wetlands. Surprisingly, gross changes towards a more natural system occurred within a relatively short timeframe. The reappearance of native vegetation, Water lilies and Bulkuru, improvements in water quality and fish biodiversity took less than 3 years. However, the weight of evidence presented here after 5 years of monitoring, shows that the abundance of native and invasive plant species appears to oscillate depending on seasonal rainfall which can induce hydrologic pressure to repress tidal water ingress, which in turn drives dissolved oxygen and temperature regimes (through vegetation and depth changes) within the wetland, effecting fish occupancy. These changes are modelled conceptually in Fig 7, describing what is happening on the Mungalla wetland post-bund removal, showing the oscillation or cycle between freshwater and saltwater tolerant plant species, associated water quality, and fish presence, concomitant with the preceding years’ weather conditions (primarily summer rainfall).

**Fig 7.**
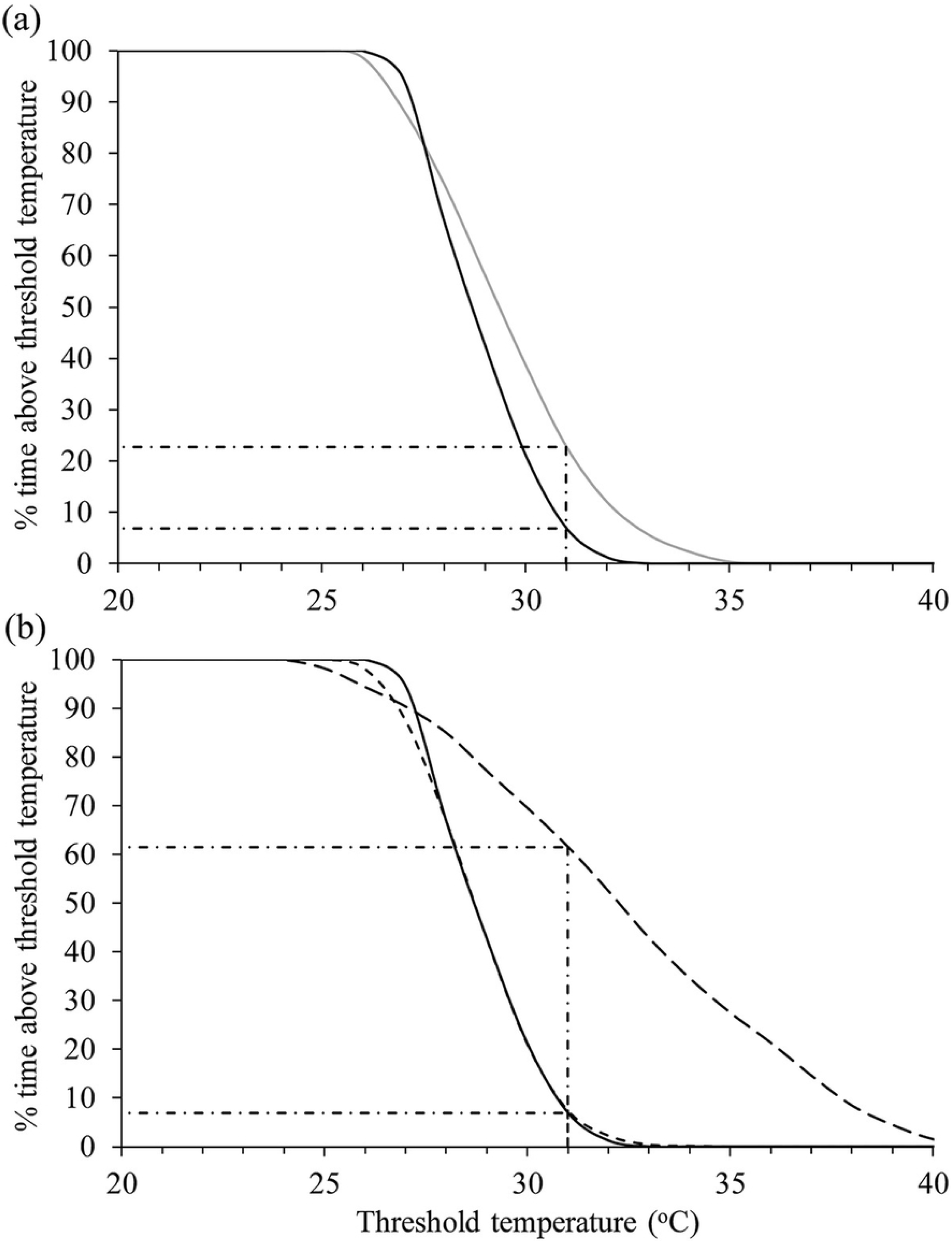
Conceptual model showing changes due to seasonal oscillations within the shallow Boolgooroo region of the Mungalla wetlands. This oscillation/cycle is likely to occur on other shallow wetlands if this method of passive remediation is employed.

Variation in wetland salinity and subsequent species and water quality variation may be an effect in the shorter term with the wetland reaching an equilibrium similar to other local natural estuarine wetlands in the longer term, i.e. Bulkuru dominant toward the seaward end of the wetlands along with other saltmarsh adapted species. Vegetation in the upper reaches will most likely remain primarily freshwater adapted species, with tidal influence in only very dry years (such as 2015). A particularly interesting outcome from this research has been the replacement of other ponded pastures by Aleman grass *(Echinochloa polystachya).* Aleman grass appears to be tolerant of marginally brackish conditions – giving it the ability to reinvade on years of lower tidal ingress, and has already shown some herbicide resistance [79]. Further research is necessary to understand more around the effects of saltwater impact on Aleman grass, including a combination of herbicide and seawater treatment, as removal of this grass may become the next challenge.

Whilst reinstatement of tidal flow has been successfully applied elsewhere to restore ecological function, this study appears to be the first of its kind targeting wetland weeds and specifically ponded pastures in the Great Barrier Reef region, and as such is an important case study for similar restoration efforts needed to effect reef water quality and the Australian government’s plan of coastal wetland restoration and protection under the Reef 2050 plan[83].

## Acknowledgements

The authors would like to acknowledge and thank the Traditional Owners, the Nywaigi people, for allowing this important project to be sited on Mungalla and for having the desire to rehabilitate their country. A special thanks to Mr J. Cassady, who encouraged and supported this project, we also acknowledge the important role Dr T. Grice has had in the establishment and conduct of the scientific monitoring. Thanks to Mr B. Butler for discussions on thermal and hypoxia tolerances on wetland aquatic species.

